# Temporal trait plasticity and neighborhood crowding predict the growth of tropical trees

**DOI:** 10.1101/2020.09.11.292615

**Authors:** Mengesha Asefa, Xiaoyang Song, Min Cao, Jesse R. Lasky, Jie Yang

## Abstract

Functional traits and neighborhood composition have been used to predict tree growth dynamics. Temporal changes in trait values (temporal trait plasticity) is one of the mechanisms for adaptive plastic response to environmental change. However, the consequence of temporal change in trait values and neighborhoods on the growth performance of individuals has rarely been investigated. We, therefore tested the effect of temporal changes in trait values and neighborhood crowding on the growth rate of individuals in a tropical forest using a dataset containing individual level growth and functional trait data for *Ficus* individuals. We collected trait and size data at two time points (2010 and 2017) for 472 individuals of 15 *Ficus* species in Xishuangbanna tropical forest dynamics plot, southwest China. We used linear mixed effect model to predict the effect of temporal trait plasticity and neighborhood crowding on the relative growth rate of individuals using these data. We found significant temporal changes in individuals’ functional traits suggesting a shift in ecological strategies from being functionally acquisitive to conservative. We also found differences in neighborhood crowding between the two census years indicating that the strength of individual interactions might change over time. The temporal changes in trait values and neighborhood crowding were found to predict better the relative growth rate of individuals, compared to static trait or crowding values in the initial and final censuses. We also found major axes of tree functional strategies in a principal component analysis, highlighting potentially adaptive trait differences. Our results in general highlight to consider the temporal dimension of functional traits and biotic interactions, as our result suggest that growth-trait relationships may vary between time points, allowing us to understand the demographic response of species to temporal environmental change.

## INTRODUCTION

Biotic interactions and environmental heterogeneity overlap spatially and temporally in effects on community assembly, creating dynamic and ecologically complex tropical forest community (Wright 2002; Zambrano et al. 2017). Biotic interactions at local scales partly drive the demographic pattern of species that ultimately shape tree community assembly (Fortunel et al. 2018). Heterogenous abiotic environments also sort species based on species ecological requirements, and regulate species performance and community dynamics in diverse systems (Lasky et al. 2013). Functional traits have been widely used to make inferences about community dynamics, as traits are believed to provide insights into the role of environment in assembly (Poorter and Bongers 2006; Yang et al. 2018). Although testing trait-growth relationships has become more common and fundamental to understand community dynamics (Swenson et al. 2017), a remaining goal is to understand this relationship from a temporal perspective.

In practice, community trait data are often collected only at a single time point. These static data are then used for downstream analyses to species demography and to understand community change through time and space (Swenson et al. 2017). While collecting a single timepoint of trait data may be pragmatic, particularly in diverse systems, a vast evolutionary ecology literature shows how traits influence individual performance (Lande and Arnold 1983, Wade and Kalisz 1990), and how traits change in adaptive (Moran 1992, Baythavong 2011), and maladaptive response to environment (Ghalambor et al. 2007). Furthermore, traditional implementation of correlating community variation in traits to demographic traits may lead to weak or mislead models and inferences (Umaña et al. 2018). How trait values and neighborhood interactions change over time, and how their temporal changes impact trees demography have not been widely tested, despite the fact that species interactions and growth strategies are known to be temporally dynamic. This may reflect the limitation of trait-based ecology as it usually gives static trait values for individuals that ignores the temporal variation of traits, and this limits the ability to understand how temporal variability in traits and biotic interactions regulates the performance of individuals over time (Swenson et al. 2017).

Adaptive phenotypic plasticity, i.e. when trait changes increase fitness, is a key strategy by which organisms respond to changes in their environment (Pigliucci 2001). Maladaptive plasticity, which is a symptom of a failure of an organism to maintain homeostasis, could also be resulted when a change in trait values through time reduce fitness of organisms (Ghalambor et al. 2007). Temporal trait plasticity is expected to increase as conditions vary over time (Lázaro-Nogal et al. 2015). Long-lived organisms have to have some level of adaptive plasticity to survive and persist through such a wide range of conditions over their lifespan, relative to short lifespan organisms such as annual plants. While a growing evolutionary ecology literature has tested for the effects of trait plasticity on intraspecific fitness variation (Dudley and Schmitt 1996, Van Kleunen and Fischer 2005), less is known about how temporal trait plasticity influences community assembly. Much of the earlier trait-based studies have focused on assessing forest community dynamics using traits measured once in the life span of trees that lacks the temporal domain of ecology (e.g., Wright et al. 2010, Lasky et al. 2014a, Paine et al. 2015, Visser et al. 2016). Temporal trait variation of communities has been less studied than temporal shifts in species composition, though traits are known to be temporally dynamic (Enquist and Enquist 2011, Fauset et al. 2012). Few studies have characterized the temporal trait changes and associated demographic consequences at community level. Van Der Sande et al. (2016) reported that community trait values (wood density increased; specific leaf area decreased) changed over time in all of the five studied forest types in the Neotropics suggesting that species shifted from being fast-growing to slow-growing species. Temporal shifts of trait distributions, mainly at the community level, have also been reported (Lasky et al. 2014b, Katabuchi et al. 2017). The decrease of specific leaf area (SLA) and leaf phosphorus content over time in the wet tropical forests also suggested a change in functional strategies of species (Muscarella et al. 2017). A long-term shift of species’ mean trait values through time showing directional change was also found in a tropical dry forest (Swenson et al. 2020). There are many studies that have inferred functional turnover in forests through time, using traits measured at a single time point and assumed to be the same for all individuals of a species. However, these studies did not measure traits of individual plants, meaning that trait plasticity could not be quantified. Testing whether the effect of traits on tree growth differs between time points and how the temporal shift of traits (i.e. temporal trait plasticity) plays a role in shaping the growth dynamics of communities may help to better understand the direction of forest structural and functional change.

Trait-growth relationships have been used to reveal plant growth strategies and predict the demographic trajectories of species (Adler et al. 2013, Yang et al. 2020 in press). However, the predictive power of traits has been sometimes weak which raises a question about the significance of traits (Paine et al. 2015). One reason for this could be, apart from the trade-offs between demographic rates that may conceal the effect of traits on species performance (Laughlin et al. 2020 in press), trait-growth relationships usually are computed at the species level using mean trait values and mean growth rates of individuals, despite the fact that trait-driven resource competition occurs at the individual level (Liu et al. 2016). Averaging trait values of individuals across the species ignores individual level trait variation, limiting the ability of traits to predict individual growth rates (Liu et al. 2016, Umaña et al. 2018). Individuals traits may predict better the growth performance of individuals, as trait differences determine individuals’ growth strategies (Yang et al. 2018, Worthy & Swenson 2019).

Neighborhood interactions influence tree growth, and can promote species diversity (Lasky et al. 2014b, Chen et al. 2016, Lamanna et al. 2017, Zambrano et al. 2017, Fortunel et al. 2018). The growth rate of individuals depends on the density of immediate neighbors with positive or negative effects. High density of neighbors often reduces the growth or survival rate of trees (Comita et al. 2010, Johnson et al. 2017, Lamanna et al. 2017). However, studies of neighborhood interactions have rarely considered temporal dynamics in biotic interactions. That is, how do neighborhood interactions change over time, and do these changes affect individual vital rates? Changes in neighborhoods over time, if overlooked, might obscure the effects of neighbors on individual growth (Bachelot et al. 2015). The number and identity of neighbors could change through time due to recruitment and mortality, and as a result the strength of neighborhood effect on growth may change over time (Newbery and Stoll 2013). One of the challenges of using neighborhood crowding covariates is that neighborhoods may change spatially in response to variation in resources (light, water, nutrients), so that the actual available resource supply might differ from what we expect from the level of crowding. And so, it may be that neighborhood dynamics are better at capturing the variation in actual resource availability, because we might expect an increase in crowding over time actually does correspond to less available resources to individuals. Thus, the effect of neighbors may not be captured unless changes in local neighborhoods are considered. However, the temporal change in neighbors and its subsequent effect on tree demography has not been widely studied, though few studies being reported.

We tested how changes in functional traits and neighborhood interactions affect the growth of species in the diverse genus of *Ficus* trees in a tropical forest. We asked the following specific questions: (i) How do traits, neighborhood crowding, and growth rate of individual trees change over time? (ii) Are functional traits and neighborhood crowding temporally consistent in predicting the relative growth rate of individuals? (iii) Does temporal trait plasticity and changes in neighborhood crowding predict better the relative growth rate of individuals compared to using only a single snapshot of traits and neighborhood crowding?

## METHODS

### Study site

We carried out this study in the 20-ha Xishuangbanna seasonal tropical rainforest dynamics plot (FDP) in southwest China (21°37′08″ N, 101°35′07″ E) (Figure S1). Dry and rainy seasons are typical features of the region with mean annual rainfall and temperature of 1493 mm and 21.8°C respectively (Cao et al. 2006). The plot ranges from 709 to 869 m in elevation (Lan et al. 2009). In 2007, all free-standing woody stems ≥ 1 cm in diameter at 130 cm from the ground (Diameter at Breast Height, DBH) were measured, mapped and identified to species (Condit 1998).

### Focal species

We used the *Ficus* (Moraceae) genus as a case study, as it is a pantropical genus with more than 800 species in the lowland tropical forest and contains functionally diverse species (Harrison 2005). *Ficus* assemblages provides a useful system to investigate the mechanisms that maintain high tropical species diversity (Lasky et al. 2014a). Furthermore, *Ficus* is the most speciose genus in the 20-ha plot, with 15 identified species and 4.6% of the total basal area in the plot, and a large quantity of soil seedbank (Tang et al. 2006). Most of the individuals are distributed on the steep slopes of the plot, and some of them are limited to ridges and valleys (Hu et al. 2012). In 2010, leaf functional traits were measured on *Ficus* individuals with a DBH of at least 10 cm with leaves accessible with pole shears (Lasky et al. 2014a) and then re-measured these trees in 2017. Thus, we used trait data for the *Ficus* trees separated by seven years and DBH data separated by ten years interval in the Xishuangbanna FDP. A species list is given in Table S1.

### Functional traits

We measured eight functional traits data in two census years for the 472 individuals of the 15 *Ficus* species in the plot. We collected five matured, healthy and sun exposed leaves for each individual in each census year and measured traits following the standardized protocols (Cornelissen et al. 2003). We measured leaf area (cm^2^), specific leaf area (cm^2^.g^−1^), leaf chlorophyll content, leaf fresh mass (g), leaf dry mass (g), leaf dry mass content, leaf thickness (mm), and leaf succulence (g.cm^-2^). These traits are expected to represent the fundamental ecological strategies of individuals for resource acquisition. Leaf area is related to light capture and heat balance (Poorter and Rozendaal 2008). Specific leaf area is linked to light interception efficiency and the main part of leaf economic spectrum (Wright et al. 2004). Leaf chlorophyll content is related to light harvesting capacity of the plant (Coste et al. 2010). Leaf thickness is related to the mechanical strength of the leaf (Onoda et al. 2011). Leaf dry matter content is associated with leaf defense ability and decomposition (Van Der Sande et al. 2016). Leaf succulence represents the trade-off of productivity and life span of the leaf (Garnier and Laurent 1994). We measured the leaf chlorophyll content using SPAD-502 Chl meter (Minolta Camera Co., Osaka, Japan), and three readings were taken at the widest portion of the leaf blades (Marenco et al. 2009). We used electronic digital micrometer to measure leaf thickness (mm) at the center of fresh leaves with multiple readings, and average was taken (Seelig et al. 2012).

### Tree growth

All *Ficus* individuals’ diameter at breast height (DBH) was measured in the Xishuangbanna FDP in 2007 and 2017. Relative growth rates (RGR) used in this study were calculated as 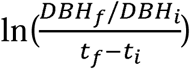, where *t* is year and the subscripts of *f* and *i* are respectively the final and initial values of the diameter at breast height (Wright et al. 2010). The relative growth rate of species is graphically indicated in Figure S2.

### Neighborhood crowding

Neighborhood competition is one of the biotic driving forces that largely determines the growth performance of individuals at the local scale (Fortunel et al. 2018). The effect of neighborhood crowding is expected to decline with distance increases from the focal stem (Uriarte et al. 2010). Here, we calculated the neighborhood competition of trees using the neighborhood crowding index (NCI) separately for the two census years in order to evaluate its temporal effect on tree growth. We computed the neighborhood crowding index (NCI) for each focal stem *i* of species *s* based on the size (DBH) and distance (d) of its neighbors (*j=1…J*) within a 15 m radius for each census year (t) (Lasky et al. 2014b, Uriarte et al. 2016). We excluded focal stems within 15 m of plot boundaries to avoid edge effects in our analysis. A 15 m radius was chosen following the previous work (Yang et al. 2020 in press).

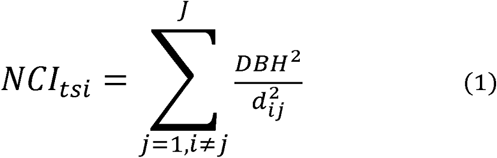

### Statistical analyses

#### The temporal shift in traits and neighborhood crowding

We first tested whether there were temporal shifts in traits (i.e. plasticity) and neighborhood crowding over time. We used a linear mixed model to test whether significant changes in univariate traits and *NCI* values occurred between the two censuses, with census as fixed effect and the variable of interest, and species identity as random factor. Furthermore, we also used the principal component analysis (PCA) on the mean centered and standardized trait values (by dividing the centered trait values by their standard deviations) to find major axes of trait variation and trait plasticity using the two censuses data.

Since functional traits were sampled twice over time on the same individuals, we were able to compare the magnitude of trait variation explained by different sources. Using traits as response variables, we included leaves, individuals, species, and census interval as random variables in our mixed-effect models to decompose and estimate the variance explained by each random variable, and expressed it in percentage as the total variance explained by each of the random components. We standardized all parameters by subtracting the minimum value from each observed value and then divided by its range value for the ease of interpretation and comparisons. Data transformation was done for all functional traits, and other variables to meet the assumption of normal distribution before analysis. Pearson correlation was carried out to check for trait covariation and hence we removed leaf fresh mass from analysis as it strongly correlated to leaf dry mass (Table S2, S3, and S4).

#### Effect of functional traits and neighborhood crowding on tree growth

The second objective of this study was to evaluate the relative importance of each functional trait and neighborhood crowding on the relative growth rate of individuals. To address this question, we built three different models: one model using the first census data, second model using the second census data, third model is using the temporal changes in trait and neighborhood crowding (i.e. the difference of traits, and neighborhood crowding between the first and second census data). For each model, we fitted individual RGR as a function of traits and neighborhood crowding using linear mixed-effects model. To handle model complexity, we fit separately the growth model for each functional trait. The first two models take the following form:

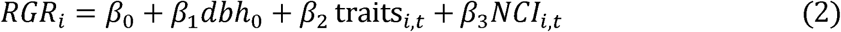

Where *dbh*_0_ is the initial tree size (DBH) at the first census year, traits _*i,t*_, represents the trait values of individual *i* in year t. *NCI*_*i,t*_, represents the *NCI* values of individual *i* in year t.β_0_ is the intercept for all individuals. For the third model, though it is the same in form with the above model, we took the temporal difference in traits and *NCI* values. We subtracted the traits/NCI values in 2010 from the corresponding values in 2017 (trait/NCI values in 2017 − trait/NCI values in 2010), and were used to describe temporal changes in traits/NCI values. We selected among models using Akaike Information Criterion (AIC) (Table S5, S6, and S7).

Additionally, we used piecewise structural equation models (SEMs) to determine any possible pathways by which traits, neighborhood crowding and initial DBH size could interactively influence the relative growth rate of individuals. We hypothesized that initial DBH size and neighborhood crowding affect individuals’ growth indirectly through their effects on functional traits. Also, the DBH size may determine the canopy position and crowding conditions of trees which may in turn influence trait expressions and ultimately affect growth of individuals. We computed a series of piecewise SEMs separately for each census data (i.e. 1^st^ census data, 2^nd^ census data, and temporal changes in traits and neighborhood crowding data). We developed a conceptual framework model that shows possible direct and indirect casual relationships among predictors and response variable (Figure S3). These hypothesized relationships help to optimize the piecewise SEMs. Functional traits, initial DBH size and neighborhood crowding were predictor variables, whereas relative growth rate of individuals was a response variable. Species were taken as random effects in our piecewise SEMs analysis. To minimize model complexity, functional traits were reduced using PCA and we used the first two PCA axes representing traits as predictors. A series of piecewise SEMs were fit to the data, and insignificant pathways were removed progressively from models to improve fitness of the model. We used Fishers’s C statistics to evaluate the goodness fit of the models with high P-values showing good fit (Lefcheck 2016). We used AIC to select the best fit and parsimonious model.

We used R version 3.5.3 to run all the analyses. ‘lme4’ package was used to fit linear mixed-effect models (Pinheiro and Bates. 2016). Principal component analysis was conducted with the ‘rda’ function in vegan package (Oksanen et al. 2014). We used ‘psem’ function in ‘piecewiseSEM’ package for piecewise SEMs analysis (Lefcheck 2016).

## RESULTS

### Temporal shifts in trait values, growth and neighborhood crowding

We tested the extent of trait, growth and neighborhood variation at the individual level and temporal time points in a tropical forest. We found significant temporal changes in trait values for at least half of the functional traits being tested (Figure 1; see also Figure S4 that compares individual traits on the scatter plot). SLA decreased significantly from the first census to the second census, whereas leaf chlorophyll content, leaf dry mass and leaf succulence increased from the first to the second census. However, we did not find significant temporal changes in trait values for Leaf fresh mass, Leaf thickness, Leaf area, and LDMC. Individuals’ size, expressed as DBH, also increased significantly over time indicating significant growth of focal trees, whereas significant change was not observed on the neighborhood crowding of individuals, consistent with the late-successional stage of the forest.

**Figure 1.**
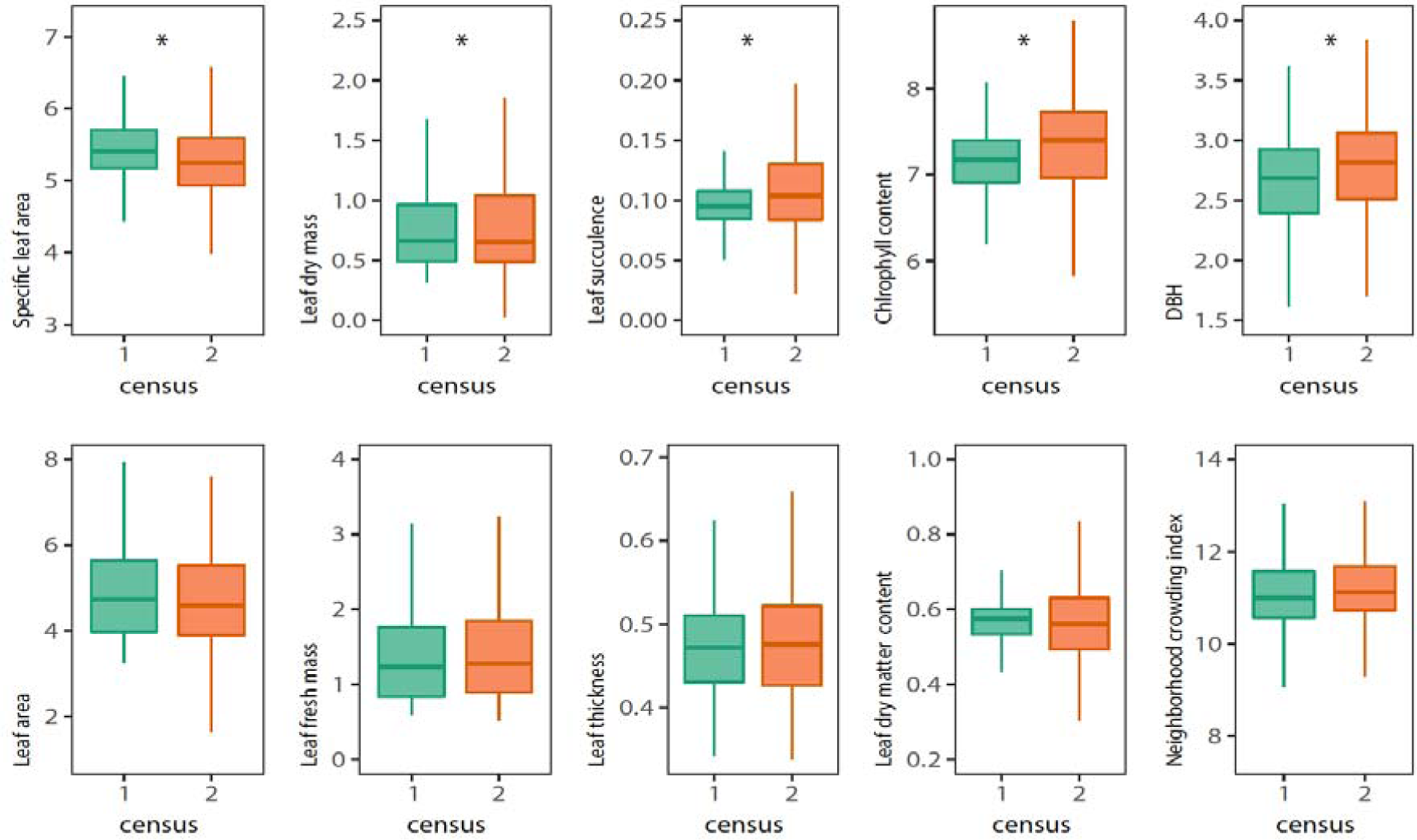
Comparison of trait values, growth and neighborhood crowding between the two census years. Asterisk (*) indicates significant differences between the censuses for each functional trait. DBH-Diameter at breast height.

We also analyzed the amount of trait variation explained by the species, individuals, years and leaves. We found that most functional traits showed significant variation among leaves, individuals, species and between census years (Figure 2). Most functional trait variations are explained by the species followed by the individual level.

**Figure 2.**
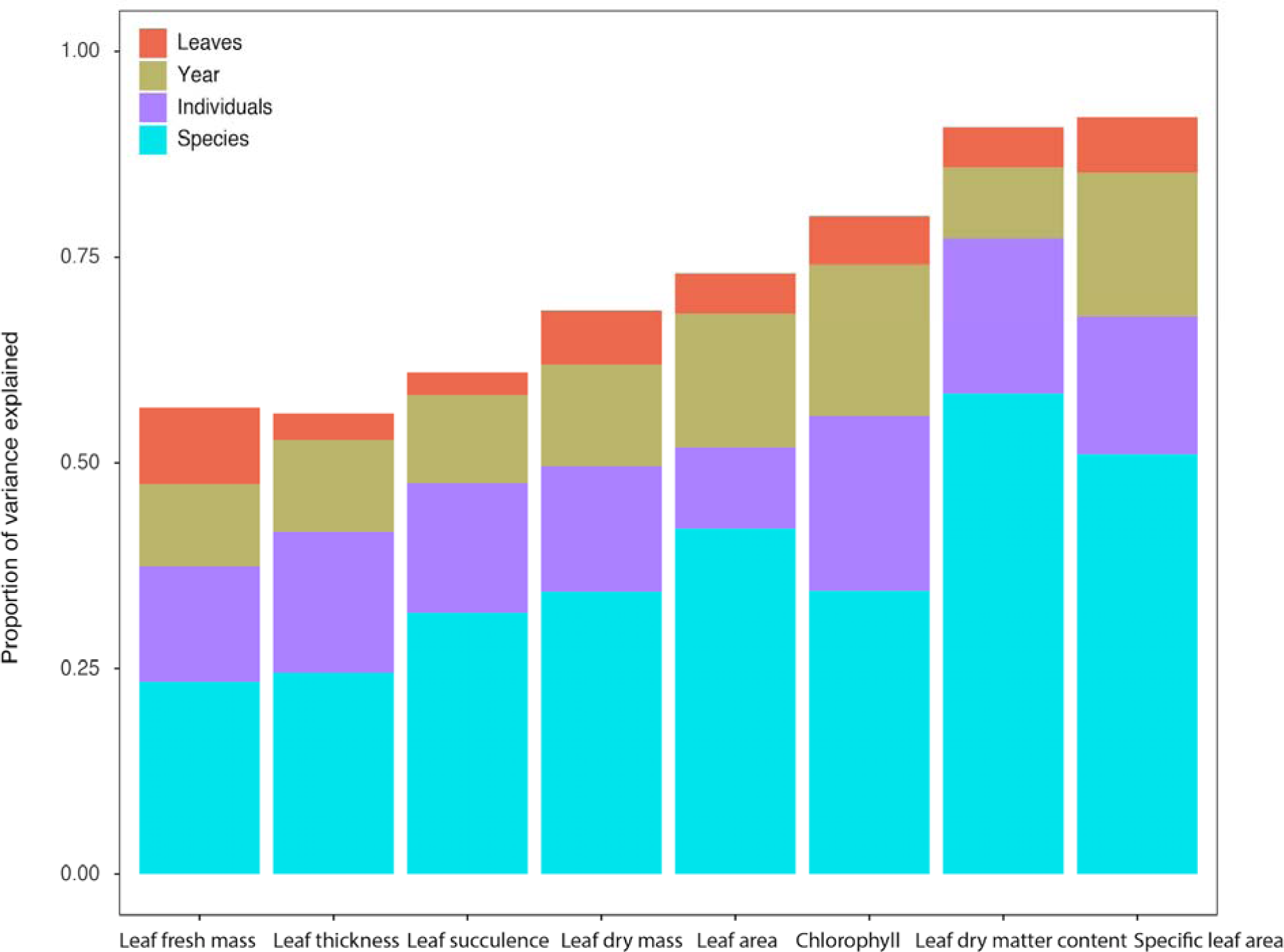
Variance in trait values explained by species, individuals, census interval, and leaves.

### Axes of functional variation

To evaluate trait associations and plant strategies, we used a PCA of the seven traits of species (Figure 3). The first two PCA axes explained almost 66 % of the variation and showed a spectrum of trait variation. The first PCA axis shows species with a large leaf area, leaf thickness and dry mass at the left to species with high SLA at the right. The second axis represents species with high chlorophyll content at the top to species with high LDMC and succulence at the bottom. These axes, therefore, represent the leaf trait spectrum tradeoff. We also conducted the PCA on the temporal change in traits (trait values in 2010 were subtracted from traits in 2017) to see axes of temporal plasticity (Figure S5). Along the first axis, individuals were separated between those species with decreasing SLA and leaf area on the right-hand side and those with increasing leaf dry mass on the left side. Species with high leaf thickness and succulence were represented at the top of the second axis.

**Figure 3.**
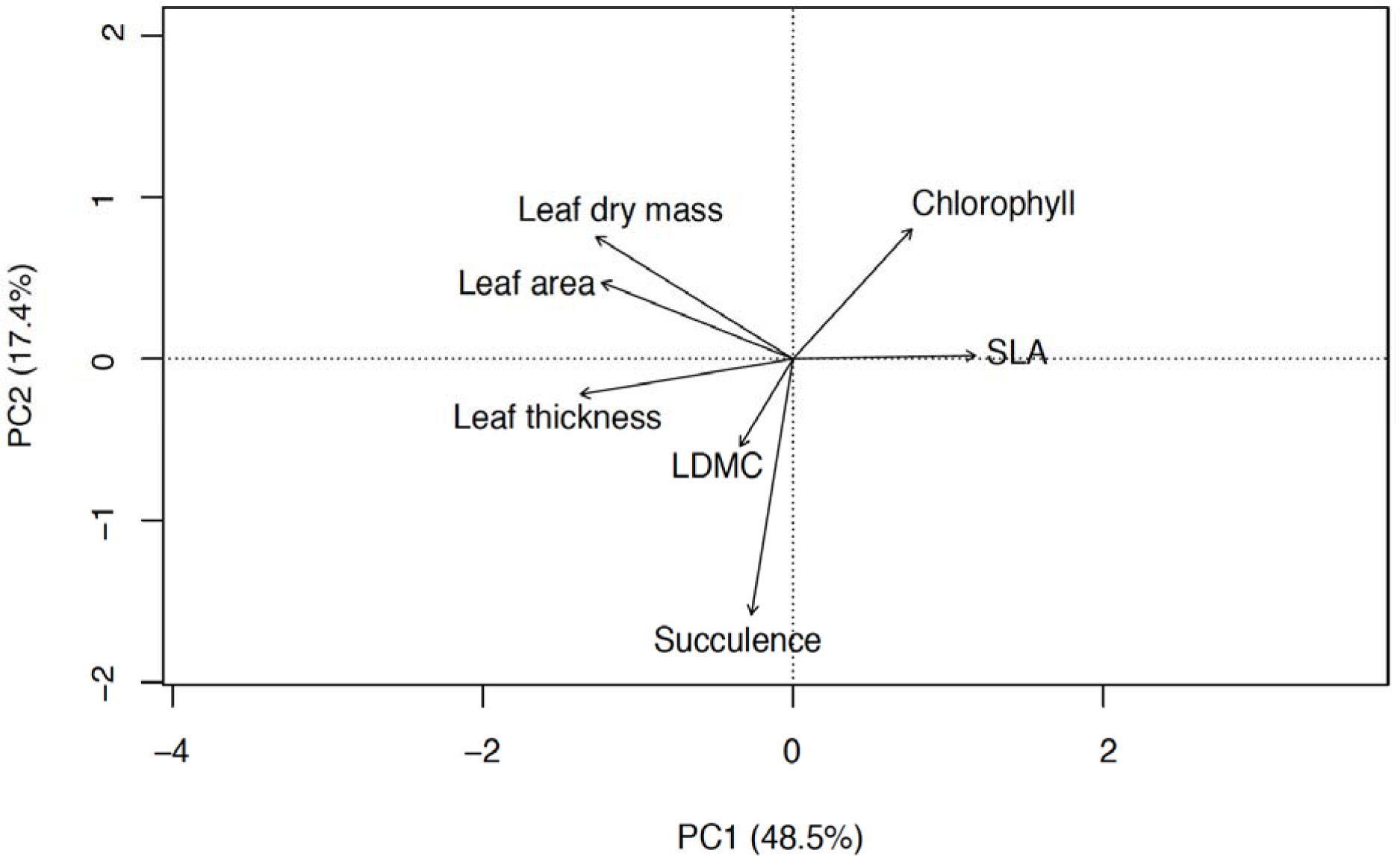
PCA representing multivariate associations among functional traits of the 472 *Ficus* individuals. The numbers in parentheses in the PC1 and PC2 axes are the variances explained by each axis.

### Effect of traits and neighborhood crowding on the relative growth rate of individual trees

We tested how growth was correlated with temporal variation of traits and neighborhood crowding. We found that traits and neighborhood crowding have not explained significantly the relative growth rate of individual trees in the first census (Figure 4a). However, in the second census leaf chlorophyll content, leaf area, leaf dry mass, LDMC, leaf succulence and neighborhood crowding significantly explained the relative growth rate of individuals (Figure 4b). We also used the temporal changes in functional traits and neighborhood crowding to predict the growth rate of individuals, and interestingly found that almost all of the change in trait values and neighborhood crowding explained better the relative growth rate of the individuals (Figure 4c). See Table S5, S6, and S7 for model AIC values.

**Figure 4.**
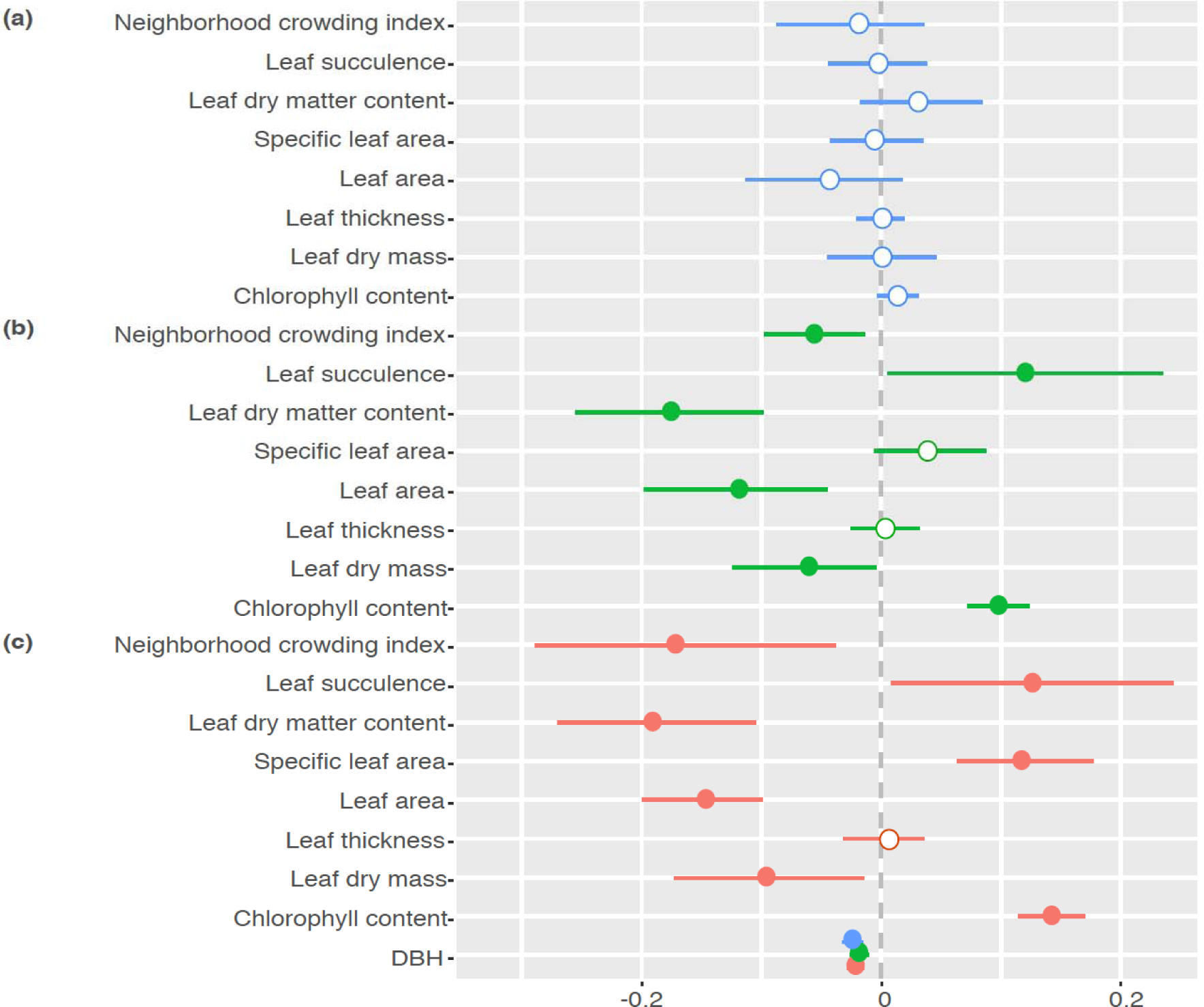
Standardized regression coefficients modelling initial size effects, traits and neighborhood effects on tree relative growth rate. (a) the first census of traits and neighborhood effect; (b) the second census of traits and neighborhood effect; (c) the effect of the change in traits and neighborhood crowding values (the trait values in 2010 were subtracted from traits in 2017) on the relative growth rate of individuals during seven years. Circles indicate posterior medians for each studied parameter and lines indicate 95% confidence intervals, with filled circles representing significant effect.

We also used SEMs to investigate any possible pathways by which traits and neighborhood crowding has interactively predicted growth of individuals. We found no significant causal relationships among traits, neighborhood crowding and initial DBH size to determine individuals’ growth rates (Figure S6). However, initial DBH size (in addition to its direct significant negative effect on RGR) has indirect significant positive effect on the RGR of individuals through its negative effect on PC1 and PC2 of the second census and temporal change data (Figure S6 b & c). We also found initial DBH to negatively be interacted with neighborhood crowding which in turn negatively influenced the RGR of individuals in the temporal change data (Figure S6 c). However, we did not find a pathway through which neighborhood crowding and traits interactively affect individuals’ growth in all census data.

## DISCUSSION

While past studies of community functional dynamics have focused on turnover in species identity, here we show patterns arising due to temporal trait plasticity of long-lived individuals. We predicted individual tree growth using traits measured on individuals while considering at the same time the biotic context in which that individual was found across time points. We showed trait-growth relationships, and negative effects of neighborhood crowding on the growth rate of individuals. The significant change of traits over time (temporal trait plasticity) and the association of functional traits with the leaf economics spectrum was also detected. Half of the functional traits measured changed significantly over time and were able to predict individual growth rates. The covariation of traits also revealed the presence of, to a certain extent, a leaf economics spectrum.

### Temporal trait plasticity and neighborhood crowding

Tropical forests inhabit dynamic environments, and therefore some changes in functional strategies of trees might be adaptive. Consistently, we found significant temporal changes of some functional traits in our plot that could potentially alter individual ecological requirements. SLA decreased and leaf dry mass increased, possibly suggesting a change in strategies for resource acquisition. Similar observations have been previously reported using data on turnover in species identities and assuming fixed trait values for species. For example, in Neotropical forests changes in species composition over time shifted toward conservative functional strategies, mainly due to disturbances (Van Der Sande et al. 2016). Muscarella et al. (2017) also found that communities shifted from species with resource acquisitive to conservative functional strategies in Mexico. Disturbances like tree fall and landslides have been common on the topographically steep plot, potentially influencing the tree community (Hu et al. 2012), especially *Ficus* species which tend to be found on the slopes. Furthermore, as species grow larger, larger amounts of energy could be invested to build non-photosynthetic tissue of the plant to maximize survival (King 2011). Thus, the formation of high leaf dry mass and succulent leaves over time could protect species from herbivore and pathogen attack, and provide mechanical support that maximize the life span of leaves and individual trees (Kitajima and Poorter 2010, Onoda et al. 2011, Poorter et al. 2018). The temporal development of this functional strategy could be associated with the resource distribution of the plot. The steep slopes of the plot, where most of the study *Ficus* species are distributed, have poor soil nutrients (Hu et al. 2012). These poor soils might influence the species to gradually develop more conservative traits (low SLA and leaf area, high leaf dry mass and leaf succulence) to maximize investment on structural components (minimize construction costs) and survival rate. Similarly, a long-term shift in functional composition due to species turnover (increased leaf area and SLA, decreased leaf succulence and wood density) was reported in a tropical dry forest (Swenson et al. 2020). Therefore, the observed temporal changes in trait values in our study, regardless of the direction (decreasing or increasing) reflects that the system of the forest is highly dynamic.

We also used the PCA of these traits to explore species functional strategies. The first two PCA axes explained 66% of the variation, and we found two lines of trait variation showing the ecological strategies of plants. The first axis corresponded to species with high SLA at one extreme, versus species with high leaf thickness and area at the other extreme. The second axis corresponded to species with high leaf chlorophyll content versus high leaf dry matter and succulent leaves. This resource use strategy trade-off is a common phenomenon in the tropical trees and is well documented (Wright et al. 2004, Katabuchi et al. 2012, Asefa et al. 2017). The negative correlation of SLA and leaf area might suggest that these two important traits were not integrated to determine the growth performance of the species (Poorter et al. 2018). Similarly, SLA and leaf area were found to be negatively correlated, probably indicating that costs to deploy SLA for large leaves was more expensive than small leaves (Milla and Reich 2007). The negative correlation of SLA with leaf thickness and/or positive correlation of LDMC with leaf thickness and succulence suggested that thick leaves maximize the longevity of trees by providing protection from herbivore attack, pathogens and physical damage. In summary, the functional trait variation of *Ficus* species supports the globally known leaf economics spectrum.

We also determined the factors associated with the greatest portion of functional trait variation. All traits were varied significantly among individuals within species and among different species. However, the largest extent of trait variations was mainly explained by species identity, with a range of 23.39% for leaf fresh mass to 58.49% for LDMC, suggesting that trait variation was stronger at the species level than the individual level. The species differences in traits might be enhanced by niche-driven evolutionary trait divergence among different species. Phylogenetically conserved traits might show small trait variation within species, suggesting less trait plasticity among individuals (Poorter et al. 2018). The detection of significant individual trait variation, however, in general highlights ecological difference among individuals. A previous study also indicated the variation of *Ficus* traits at the individual level, reflecting differences of ecological requirements among individuals co-occurring together at small scales (e.g. 10 m) (Lasky et al. 2014a). Our result highlights that individual trait variation supports the species level variation of functional traits suggesting that both the individual and species level approach together helps to better understand community dynamics.

### Effect of traits on the relative growth rate of individuals

We tested to what extent individual trait variation predicts the individual variation of growth rate. The relative growth rate of individuals was found to vary substantially among individuals of the same species and among different species (Figure S2). We found that initial DBH has consistent relationships with growth of individuals in both censuses. Our results indicate that the relationship between functional traits and relative growth rate varied through time. Functional traits measured at the first census did not predict the growth rates of individuals. However, in the second census, leaf dry mass, LDMC and leaf area negatively predicted individuals’ growth rates, whereas chlorophyll content and leaf succulence were positively associated with the growth rate of individuals.

Detection of weak trait−growth relationships in the first census could be attributed to different factors. Using stem tree diameter as a growth indicator might be a poor parameter to describe the entire plant growth pattern, especially for small plants due to the fact that plants could invest their energy in height and leaf growth to capture adequate amount of sunlight as height growth is more ecologically important than diameter growth, or underground investment to maximize nutrient acquisition (Paine et al. 2015, Poorter et al. 2018). The trait-growth relationship might also be confounded by developmental stages of trees, as ontogenetic stages of trees were found to determine trait-growth relationships (Iida et al. 2014, Lasky et al. 2015, Visser et al. 2016), suggesting size-dependent changes in growth strategies (Gibert et al. 2016). However, these developmental differences should be relatively subtle given the short time interval between censuses (7 years) relative to the lifespan of many trees (many decades).

The relative growth rate of individuals in the second census, however, was found to be positively correlated with SLA and chlorophyll content of the species. This is consistent with the expectation that high chlorophyll content and SLA are considered to maximize the efficiency of biomass investment for light interception (Poorter et al. 2008). Similarly, Poorter and Bongers (2006) found that SLA predicted higher growth rate of rainforest species. Lack of consistent predictive power of traits on the relative growth rate of individuals over time points in this study, however, might indicate how sensitive tropical forests are to the temporal dynamics of the environment or trait plasticity, indicating the importance of complexity and temporal dynamics in tropical rainforests. However, temporal trait plasticity did explain the relative growth rate of individuals. Increases in leaf chlorophyll content and leaf succulence over time had positive correlations with growth, while a decrease in leaf area, leaf dry mass and LDMC over time had a negative effect on growth, suggesting that temporal shifts in trait values appeared to be more predictive of growth rate than initial census trait values. These functional traits showed large variation across time points and subsequently were found to be growth determinants. Considering the effect of temporal trait plasticity helps to better predict growth dynamics of trees. Our findings in general highlight that considering the dynamic ecological dimension of species such as traits, helps to gain a temporal understanding of plant growth dynamics.

We also found both direct and indirect effects of initial DBH of trees on the growth rate of individuals from our SEMs analyses. Initial DBH of trees was found to directly negatively affect the relative growth rate of individuals, as we found in the mixed effects models. This might be because small adult size plants could allocate more resources to height than diameter growth, as height is more important for interception of light resource (Paine et al. 2015). However, initial DBH of trees indirectly positively affected the relative growth rate of trees through its significant negative effect on PC1 and PC2 in the second census and PC1 in the temporal change data. The PC1 both in the second and temporal change data were mostly represented by leaf dry mass and LDMC, while SLA and succulence were largely represented by PC2 in the second census data (Table S8). Initial DBH size of trees could be negatively related with resource conservative traits mentioned above that may provide protection against herbivores and pathogens (Van Der Sande et al. 2016). Plants could prioritize their survival by building non-photosynthetic tissues particularly at small adult size at which susceptibility to herbivory and physical damages is higher. This reduces mortality and maximizes the longevity of trees (Onoda et al. 2011), and provides time for trees to gradually shift to the strategy by which more energy could be invested for their growth rates (Iida et al. 2014). Consistently, our result may demonstrate that initial DBH size of trees could indirectly promote the growth rate of individuals in a long-term by controlling trait expressions in response to many biotic and abiotic factors. Initial DBH size of trees also negatively interacted with neighborhood crowding to significantly limit the RGR of individuals in the temporal change data. The crowding conditions of individuals could be influenced by the size of neighboring trees. Large trees might dominate small neighborhood individuals through competition thereby reducing neighborhood density and/or limit individuals’ growth, as they may have large canopy, crown and deep root systems (Yang et al. 2020 in press). Our result therefore, suggests that various factors may interactively influence species performance through multiple pathways.

### Effect of neighborhood crowding on the relative growth rate of individuals

We evaluated the effect of neighborhood interactions on individual growth, and how the effect changes over time. We found that neighborhood crowding significantly limits the growth rate of individuals, which is consistent with previous studies (Bagchi et al. 2014, Lasky et al. 2014b, Fortunel et al. 2016, Liu et al. 2016, Fortunel et al. 2018, Umaña et al. 2018). Also, species-specific negative density dependence was found to drive seedling survival (Lin et al. 2012). This further confirmed our result that species growth might be influenced by biotic interaction with neighbors. Interestingly, our result demonstrates not only the negative effect of neighborhood crowding on growth but also its effect found to vary through time.

Similarly, the growth rate of individuals was negatively affected by changes in the number of conspecifics over time (Umaña et al. 2018). We detected a coordinated shift (temporal plasticity) of species from acquisitive to conservative, which may enhance the density-dependent effect of neighbors on growth rate due to niche overlap (Uriarte et al. 2010). However, the effect of neighborhood crowding on individuals’ performance in the first census was not significant, suggesting that interactions between tree neighbors might not be consistent over time. This inconsistency may be related to trait differences among individuals, which was may be an important mechanism of coexistence (Lasky et al. 2014a). The relationship of growth and neighborhood competition might not be completely captured under current environmental dynamics. As a result, it is always a challenge to explore the effect of neighborhood competition dynamics on the tree performance unless a temporal change in local neighbor density is considered. We found that temporal changes in neighborhood crowding affected tree growth, suggesting the importance of the approach we used to test the impact of temporal shifts of neighborhood competition on the demography of species. The result of this study highlights the importance of the temporal dimension of ecology to understand better how species interactions change over time and predict individual performance.

In conclusion, our result demonstrates that functional traits and neighborhood crowding have changed over time. This temporal trait plasticity was found to best predict the growth rate of individuals. Neighborhood interactions also limited the growth of individuals. This all together suggested that the temporal dynamics of traits and biotic interactions need to be considered to explain better the growth dynamics of tropical trees. Furthermore, trees tended to shift their functional strategies from being acquisitive to conservative, as we observed the increment of leaf dry mass and succulence, and decrement of SLA and leaf area through time points. We also found major axes of tree functional strategies in PCA, highlighting potentially adaptive trait differences. This study in general highlights that a temporal-based approach of investigating the relationship between traits and biotic interactions, and growth for each individual tree, can help gain insights into forest dynamics. To better predict future changes in community structure, function and dynamics, it is therefore important to consider the temporal change of environments together with changes in traits and biotic interactions over time, as trait-neighborhood-performance relationships vary temporally with environmental conditions.

## ACKNOWLEDGMENTS

This research was supported by the Strategic Priority Research Program of the Chinese Academy of Sciences (XDB31000000), National Natural Science Foundation of China (31800353 and 31670442), the CAS 135 program (No. 2017XTBG-T01), the Chinese Academy of Sciences Youth Innovation Promotion Association (2016352), the Southeast Asia Biodiversity Research Institute, Chinese Academy of Sciences (Y4ZK111B01), the West Light Foundation of the Chinese Academy of Sciences, and the “Ten Thousand Talents Program of Yunnan” (YNWR-QNBJ-2018-309). Data collection in 2010 was supported by a U.S. National Science Foundation East Asia Pacific Summer Institute fellowship, and a Casey Stengl research fellowship from the Graduate Program in Ecology, Evolution, and Behavior at the University of Texas–Austin to JRL.

## AUTHORS’ CONTRIBUTIONS

MA, YJ and XS designed the study; YJ, JRL and XS collected the field data; MA, YJ and XS analyzed the data; MA, YJ, XS and JRL wrote the manuscript, and all authors provided comments.

## DATA AVAILABILITY STATEMENT

Data will be available online upon manuscript acceptance.

## Supporting information

Figure S1. Location map of the study area

Figure S2. Relative growth rate of species

Figure S3. Framework of the structural equation model

Figure S4. Scatter plot of functional traits

Figure S5. PCA showing axes of temporal trait plasticity

Figure S6. Piecewise structural equation model showing trait, DBH, and neighborhood relationship with growth

Table S1. List of *Ficus* species

Table S2. Pearson correlation of traits in the first census

Table S3. Pearson correlation of traits in the second census

Table S4. Pearson correlation of temporal changes in traits

Table S5. AIC values of mixed-effect models in the first census

Table S6. AIC values of mixed-effect models in the second census

Table S7. AIC values of mixed-effect models of the temporal change in traits and NCI

Table S8. Trait loads on the PCA axes

