## supplemental file for "Temporal trait plasticity and neighborhood crowding predict the growth of tropical trees"

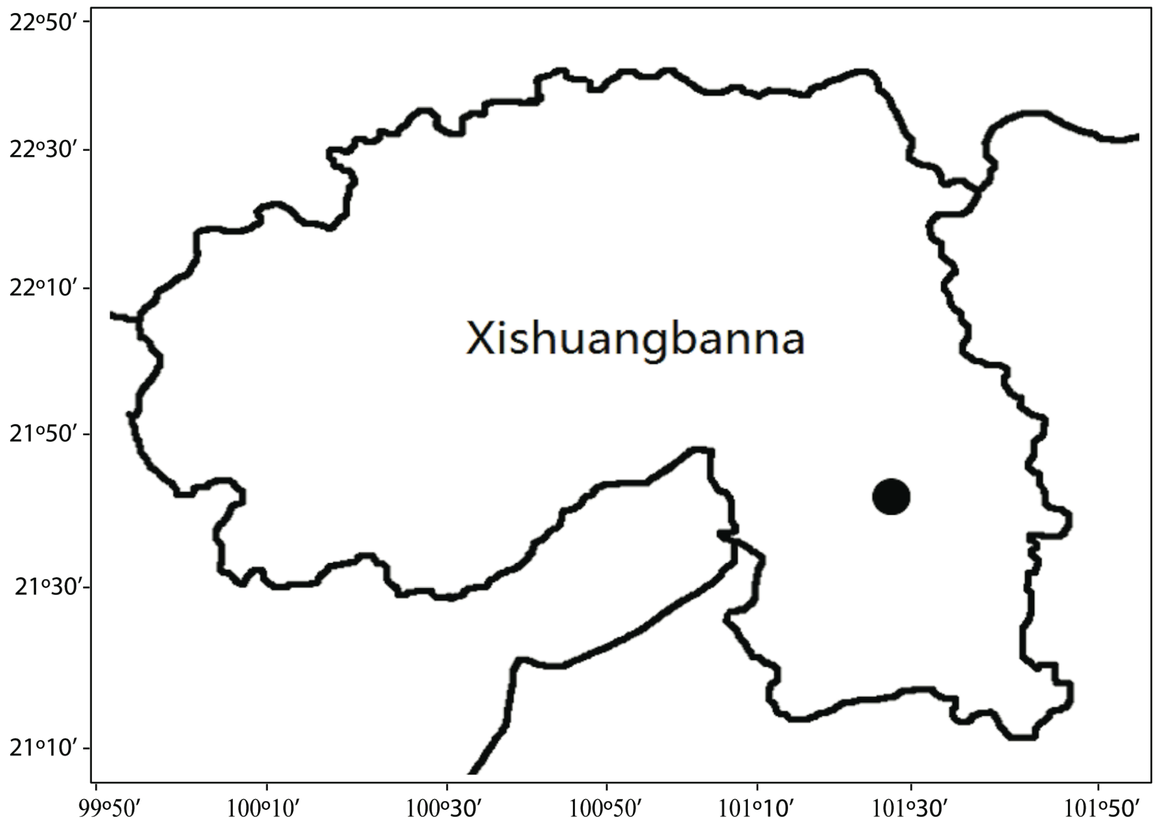


Figure S1. Location map of the 20-ha Xishuangbanna seasonal tropical forest dynamics plot in China (the black circle dot). The map was generated using ArcGIS 10.1 (www.esri.com).

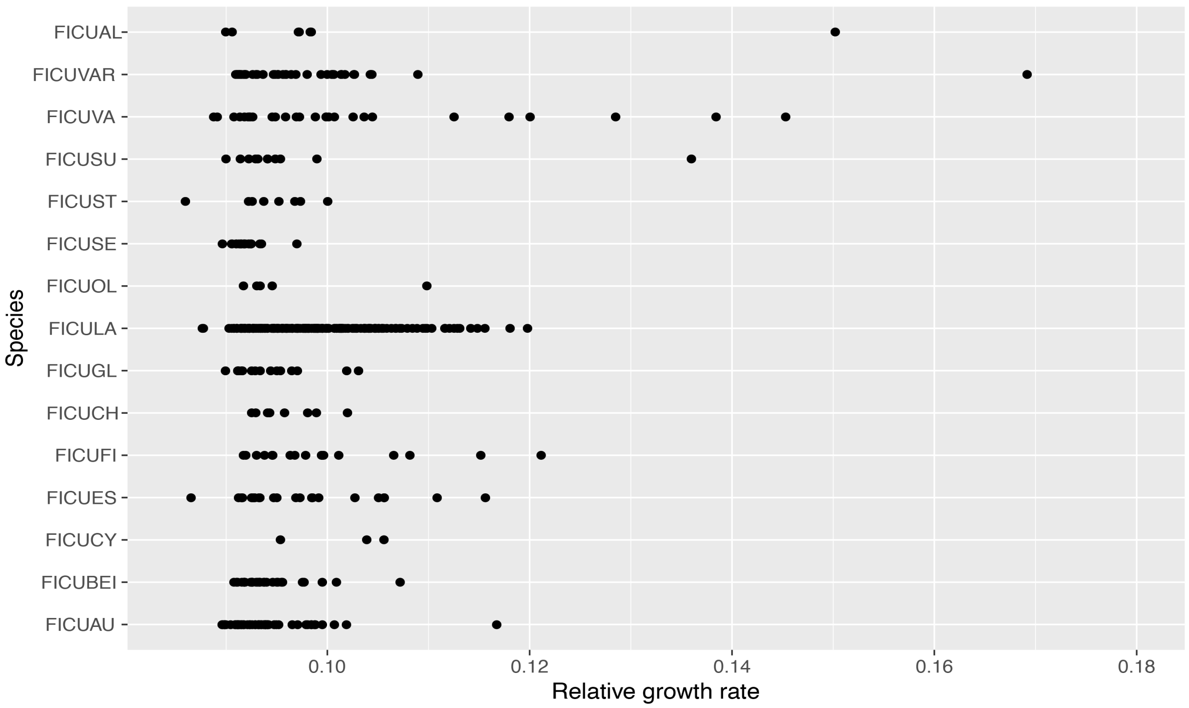


Figure S2. The relative growth rate of *Ficus* species for the 10 years data. Each point is an individual tree.

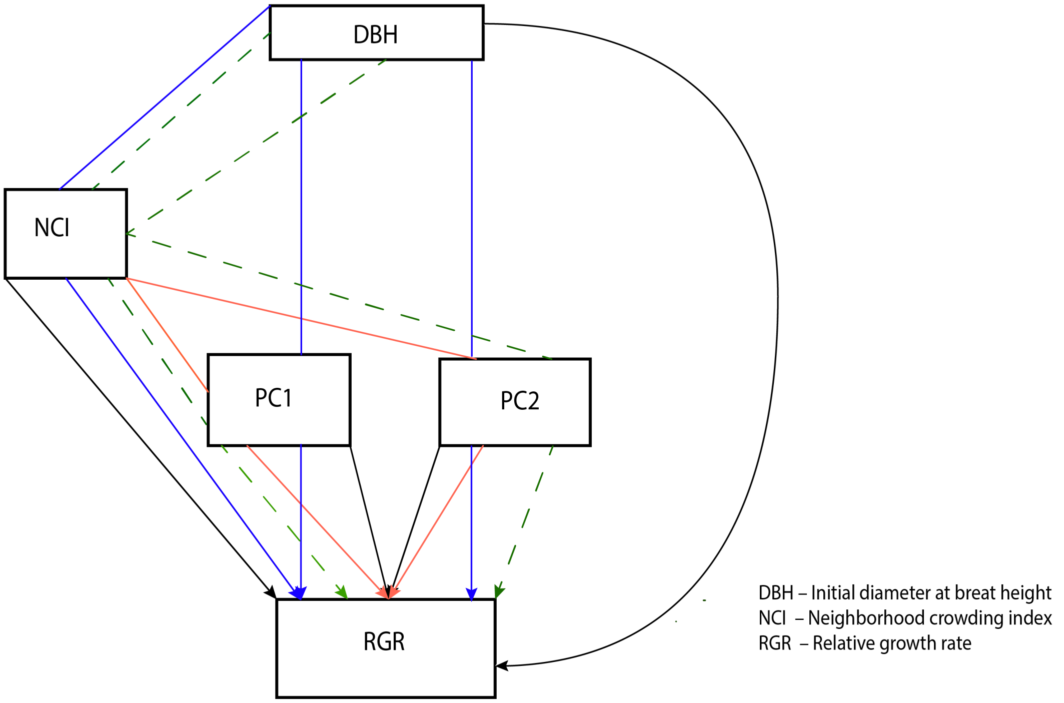


Figure S3. A Conceptual framework of the structural equation model hypothesizing the possible direct and indirect pathways of causal relationships among predictors and response variables. The black lines are the direct effect of predictors on the response variable (RGR). The blue and red lines show the causal relationships between two predictors and their effect on the RGR. The green dotted lines show the casual relationships among the three predictor variables and their consequences on the RGR. PC1 and PC2 represent functional traits from 'rda' analysis.

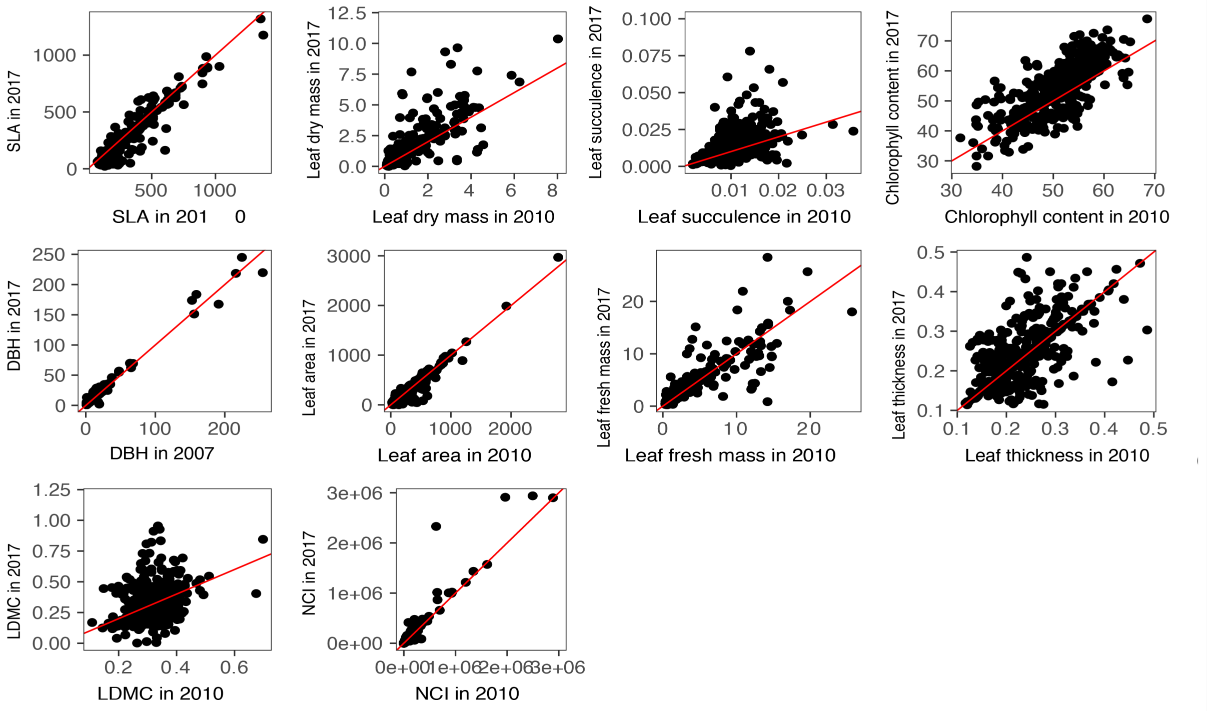


Figure S4. Scatter plot comparing trait values between the two census data using identity line with slope one (the red line). The more similar in trait values between the two censuses data, the more the individuals tend to concentrate in the vicinity of identity line.

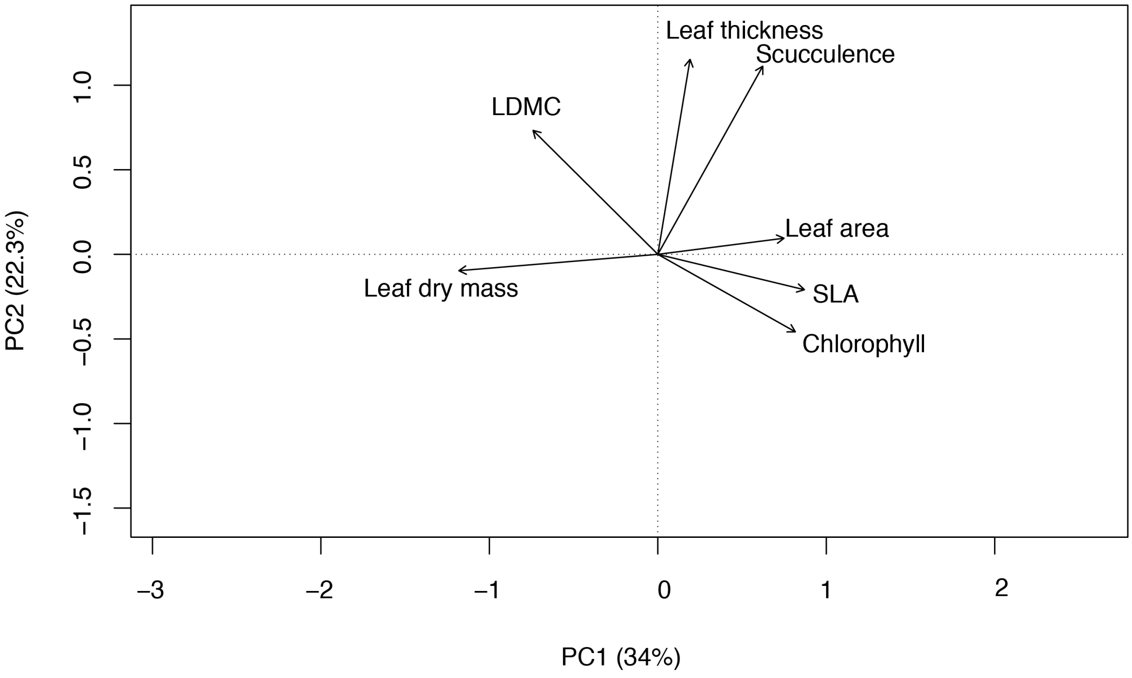


Figure S5. Principal components analysis (PCA) on the temporal change in traits showing temporal plasticity among functional traits of the 472 *Ficus* individuals. The numbers in parentheses in the PC1 and PC2 axes are the variances explained by each axis.

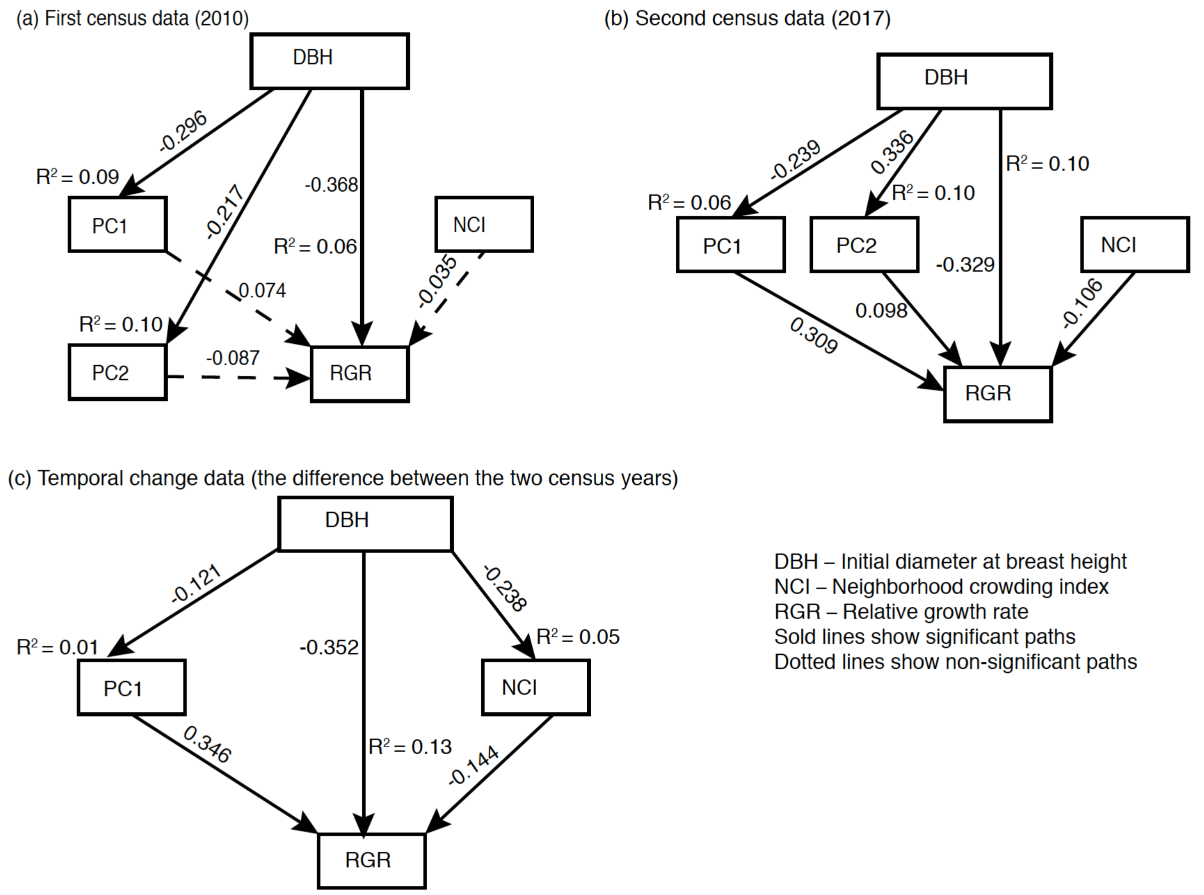


Figure S6. Piecewise structural equation model that shows pathways of the effect of functional traits, neighborhood crowding and initial DBH size of trees on the relative growth rate of individuals. Numbers on the arrows are standardized path coefficients of models. (a) AIC=76, Fisher's C=46, P-value=0, df=6; (b) AIC=30.9, Fisher's C=0.88, P-value=0.99, df=6; (c) AIC=33.8, Fisher's C=5.86, P-value=0.053, df=2.

Table S1. List of the study *Ficus* species

| | | |
|---|---|---|
| FICUAU | *Ficus* | *auriculata* |
| FICUBEI | *Ficus* | *benjamina* |
| FICUCY | *Ficus* | *cyrtophylla* |
| FICUES | *Ficus* | *esquiroliana* |
| FICUFI | *Ficus* | *fistulosa* |
| FICUCH | *Ficus* | *chrysocarpa* |
| FICUGL | *Ficus* | *glaberrima* |
| FICULA | *Ficus* | *langkokensis* |
| FICUOL | *Ficus* | *oligodon* |
| FICUSE | *Ficus* | *semicordata* |
| FICUST | *Ficus* | *stricta* |
| FICUSU | *Ficus* | *subincisa* |
| FICUVA | *Ficus* | *vasculosa* |
| FICUVAR | *Ficus* | *variolosa* |
| FICUAL | *Ficus* | *altissima* |

Table S2 Coefficients of Pearson correlation among functional traits in the 1$^{st}$ census (2010). Significant correlations are marked with bold

| | Chol | Fresh mass | Dry mass | thickness | Leaf area | SLA | LDMC | Succulence |
|---|---|---|---|---|---|---|---|---|
| Chol | 1.00 | | | | | | | |
| Fresh mass | -0.09 | 1.00 | | | | | | |
| Dry mass | -0.05 | **0.97** | 1.00 | | | | | |
| thickness | **-0.14** | **0.51** | **0.50** | 1.00 | | | | |
| Leaf area | -0.05 | **0.76** | **0.75** | **0.22** | 1.00 | | | |
| SLA | -0.03 | **-0.14** | **-0.20** | **-0.39** | **0.24** | 1.00 | | |
| LDMC | **0.36** | **-0.26** | **-0.09** | **-0.16** | **-0.25** | **-0.36** | 1.00 | |
| Succulence | **-0.19** | **0.50** | **0.46** | **0.57** | 0.02 | **-0.61** | **-0.29** | 1.00 |

Table S3. Coefficients of Pearson correlation among functional traits in the 2$^{nd}$ census (2017. Significant correlations are marked with bold

| | Chol | Fresh mass | Dry mass | thickness | Leaf area | SLA | LDMC | Succulence |
|---|---|---|---|---|---|---|---|---|
| Chol | 1.00 | | | | | | | |
| Fresh mass | **-0.20** | 1.00 | | | | | | |
| Dry mass | **-0.20** | **0.94** | 1.00 | | | | | |
| Thickness | **-0.20** | **0.40** | **0.38** | 1.00 | | | | |
| Leaf area | **-0.11** | **0.71** | **0.62** | **0.11** | 1.00 | | | |
| SLA | **0.14** | **-0.21** | **-0.27** | **-0.41** | **0.18** | 1.00 | | |
| LDMC | **-0.18** | **0.24** | **0.41** | 0.02 | **0.22** | **-0.20** | 1.00 | |
| Succulence | 0.04 | 0.00 | -0.01 | 0.04 | -0.06 | **-0.12** | -0.05 | 1.00 |

Table S4. Coefficients of Pearson correlation among temporal changes in functional traits. Significant correlations are marked with bold

| | Chol | Fresh mass | Dry mass | thickness | Leaf area | SLA | LDMC | Succulence |
|---|---|---|---|---|---|---|---|---|
| Chol | 1.00 | | | | | | | |
| Fresh mass | **-0.13** | 1.00 | | | | | | |
| Dry mass | **-0.23** | **0.69** | 1.00 | | | | | |
| thickness | -0.05 | 0.08 | 0.08 | 1.00 | | | | |
| Leaf area | **-0.30** | **0.50** | **0.37** | -0.05 | 1.00 | | | |
| SLA | **0.29** | **-0.28** | **-0.46** | -0.08 | -0.08 | 1.00 | | |
| LDMC | **-0.33** | **0.28** | **0.55** | -0.05 | **0.44** | **-0.34** | 1.00 | |
| Succulence | **0.13** | 0.04 | -0.02 | 0.02 | -0.01 | 0.03 | -0.06 | 1.00 |

Table S5. AIC values for the model effect of traits and NCI values on the relative growth rate of individuals in the first census year

| Models | AIC |
|---|---|
| $dbh_0$ | -1285.2376 |
| $dbh_0$ + NCI | -1278.8043 |
| $dbh_0$ + Chlorophyll | -1278.227 |
| $dbh_0$ + Chlorophyll + NCI | -1271.6922 |
| $dbh_0$ + dry mass | -1278.084 |
| $dbh_0$ + dry mass + NCI | -1271.6778 |
| $dbh_0$ + leaf thickness | -1277.0491 |
| $dbh_0$ + leaf thickness + NCI | -1270.6225 |
| $dbh_0$ + leaf area | -1280.044 |
| $dbh_0$ + leaf area + NCI | -1273.6411 |
| $dbh_0$ + SLA | -1277.741 |
| $dbh_0$ + SLA + NCI | -1271.2648 |
| $dbh_0$ + LDMC | -1279.3831 |
| $dbh_0$ + LDMC + NCI | -1272.8814 |
| $dbh_0$ + succulence | -1277.4209 |
| $dbh_0$ + succulence + NCI | -1271.0137 |

Table S6. AIC values for the model effect of traits and NCI values on the relative growth rate of individuals in the second census year.

| Models | AIC |
|---|---|
| $dbh_0$ | -1285.2376 |
| $dbh_0$ + NCI | -1284.3847 |
| $dbh_0$ + Chlorophyll | -1339.0258 |
| $dbh_0$ + Chlorophyll + NCI | -1338.1874 |
| $dbh_0$ + dry mass | -1282.7945 |
| $dbh_0$ + dry mass + NCI | -1282.0704 |
| $dbh_0$ + leaf thickness | -1276.8017 |
| $dbh_0$ + leaf thickness + NCI | -1275.9299 |
| $dbh_0$ + leaf area | -1290.1073 |
| $dbh_0$ + leaf area + NCI | -1289.0869 |
| $dbh_0$ + SLA | -1278.6412 |
| $dbh_0$ + SLA + NCI | -1277.8729 |
| $dbh_0$ + LDMC | -1299.3965 |
| $dbh_0$ + LDMC + NCI | -1298.8689 |
| $dbh_0$ + succulence | -1284.1208 |
| $dbh_0$ + succulence + NCI | -1283.1475 |

Table S7. AIC values for the model effect of temporal changes in traits and NCI values on the relative growth rate of individuals.

| Models | AIC |
|---|---|
| $dbh_0$ | -1285.2376 |
| $dbh_0$ + NCI | -1286.3911 |
| $dbh_0$ + Chlorophyll | -1399.1448 |
| $dbh_0$ + Chlorophyll + NCI | -1403.7686 |
| $dbh_0$ + dry mass | -1284.9278 |
| $dbh_0$ + dry mass + NCI | -1286.7761 |
| $dbh_0$ + leaf thickness | -1277.4317 |
| $dbh_0$ + leaf thickness + NCI | -1278.4659 |
| $dbh_0$ + leaf area | -1320.2941 |
| $dbh_0$ + leaf area + NCI | -1326.3303 |
| $dbh_0$ + SLA | -1297.3738 |
| $dbh_0$ + SLA + NCI | -1299.2663 |
| $dbh_0$ + LDMC | -1304.789 |
| $dbh_0$ + LDMC + NCI | -1310.0319 |
| $dbh_0$ + succulence | -1284.2068 |
| $dbh_0$ + succulence + NCI | -1285.4509 |

Table S8. Trait loads of the first two PCA axes

| Traits | 1st census (2010) | | 2nd census (2017) | | Temporal change | |
|---|---|---|---|---|---|---|
| | PC1 | PC2 | PC1 | PC2 | PC1 | PC2 |
| Chlorophyll | 0.6020 | -0.8675 | 1.11546 | -0.0405 | 1.6933 | -0.3543 |
| Dry mass | -2.3096 | 0.6160 | -2.47490 | -0.4156 | -2.1422 | 0.2579 |
| Thickness | -2.2267 | -0.4317 | -1.58352 | 1.40667 | 0.1646 | 2.2668 |
| Leaf area | -1.4108 | 1.8183 | -1.76714 | -1.6680 | -1.7812 | -1.1675 |
| SLA | 1.2981 | 2.2851 | 1.15949 | -2.1299 | 1.6316 | -1.1879 |
| LDMC | 0.7849 | -1.7778 | -1.58040 | -0.4518 | -2.2675 | -0.3640 |
| Succulence | -2.2338 | -0.8856 | 0.04845 | 1.07036 | 0.3502 | -0.5348 |
